# Inadequate sampling of the soundscape leads to overoptimistic estimates of recogniser performance: a case study of two sympatric macaw species

**DOI:** 10.1101/2022.12.29.522205

**Authors:** Thomas C. Lewis, Ignacio Gutiérrez Vargas, Andrew P Beckerman, Dylan Z. Childs

**Affiliations:** Department of Biosciences, Alfred Denny Building, University of Sheffield, Sheffield, South Yorkshire, United Kingdom, S10 2TN; University of Costa Rica; Macaw Recovery Network, Punta Islita, Nandayure, Guanacaste, Costa Rica

**Keywords:** Passive acoustic monitoring, machine learning, training dataset, iteration, species-specific

## Abstract

Passive acoustic monitoring (PAM) – the use of autonomous recording units to record ambient sound – offers the potential to dramatically increase the scale and robustness of species monitoring in rainforest ecosystems. PAM generates large volumes of data that require automated methods of target species detection. Species-specific recognisers, which often use supervised machine learning, can achieve this goal. However, they require a large training dataset of both target and non-target signals, which is time-consuming and challenging to create. Unfortunately, very little information about creating training datasets for supervised machine learning recognisers is available, especially for tropical ecosystems. Here we show an iterative approach to creating a training dataset that improved recogniser precision from 0.12 to 0.55. By sampling background noise using an initial small recogniser, we can address one of the significant challenges of training dataset creation in acoustically diverse environments. Our work demonstrates that recognisers will likely fail in real-world settings unless the training dataset size is large enough and sufficiently representative of the ambient soundscape. We outline a simple workflow that can provide users with an accessible way to create a species-specific PAM recogniser that addresses these issues for tropical rainforest environments. Our work provides important lessons for PAM practitioners wanting to develop species-specific recognisers for acoustically diverse ecosystems.

## Introduction

Effective monitoring of wildlife populations is required to mitigate rapid and widespread environmental change (Gibbs, Snell and Causton, 1999; Pereira and David Cooper, 2006; Nichols et al., 2015). Long-term, standardised monitoring provides data on the presence or abundance of target species, which is necessary to identify the factors affecting population growth, abundance, and persistence (Pollock et al. 2002; Fedy and Aldridge 2011; Nuttall et al. 2022). These data can be challenging to acquire for wide-ranging species of conservation concern, as they are often found at low density in inaccessible environments such as tropical forests or marine ecosystems (Barnes, 2001; Guschanski et al., 2009; Dénes, Tella and Beissinger, 2018). The challenge is to collect sufficient volumes of ecologically relevant data at appropriate spatial scales. However, traditional survey methods are not well-suited to meet this challenge because they are often impractical and labour-intensive.

Passive Acoustic Monitoring (PAM) has emerged as a cost-effective method to address this challenge. It is one of several advances in high-throughput sensing technologies, such as remote sensing, LIDAR, and camera traps, that can scale up data collection while maintaining or minimising work effort on the ground (Gibb et al., 2019). PAM is a rapidly expanding field, benefitting from developments and cost reductions tied to hardware such as Automated Recording Units (ARUs) (Snaddon et al., 2013; Hill et al., 2019; Teixeira, Maron and Rensburg, 2019). These developments dramatically increase the quality and quantity of ecological data collected (Gibb et al., 2019). ARUs provide an efficient and non-invasive data collection platform to inform a wide variety of ecological metrics, including community composition (Pillay et al., 2019; Bradfer-Lawrence et al., 2020), abundance (Marques et al., 2013; Pérez-Granados et al., 2019), occupancy (Wood et al., 2019) and individual breeding biology (Marin-Cudraz et al., 2019).

Like any technology, PAM engenders several practical challenges. Critically, PAM requires expertise in post-collection data processing to collect meaningful information, such as detections of target species from raw audio files. For example, users can extract data manually by labelling target signals (Campos-Cerqueira and Aide, 2016; Abrahams and Geary, 2020), though this approach is time-consuming and requires expert knowledge. Due to the large quantities of data produced, there is increasing interest in developing machine learning classifiers to automate target species identification from raw audio files. In addition, deep-learning classifiers can be highly accurate at bioacoustic tasks, though they require a large amount of data to train (Bermant et al., 2019; Stowell et al., 2019; Zhong et al., 2020). Such classifiers are increasingly used to track individual species in the wild, for example, sperm whales (*Physeter macrocephalus*; Bermant et al., 2019) and the Northern grey gibbon (*Hylobates funereus;* Clink and Klinck, 2020). More generalised deep-learning classifiers to identify multiple species have also been developed, though currently, most are insufficient to classify all species of interest in many ecological applications (Ventura et al., 2015; Stowell et al., 2019; Zhong et al., 2020). BirdNet was the first generalised bioacoustic classifier for avian species. This deep artificial neural network (DNN) can identify 984 North American and European bird species with a mean precision of 79% in the presence of background noise representing a considerable step forward in the generalised classification (Kalh et al. 2021).

Developing effective classifiers for use with PAM can be time-consuming and prohibitively complex for non-expert users (Gibb et al., 2019). Proprietary software such as Kaleidoscope (Wildlife Acoustics, USA) and cloud-based platforms such as Arbimon (Aide et al., 2013; Bravo, Berríos and Aide, 2017) offer a potential solution. However, such tools are often limited in their options for species identification tasks and, in some cases, incur a prohibitive cost for many conservation projects. Thus, where relevant off-the-shelf classifiers or suitable platforms are unavailable, custom classifiers must still be developed for individual applications on a case-by-case basis. These domain-specific classifiers are often created using simple machine learning methods (sML). These have a lower technical barrier to entry than deep-learning classifiers and can be more suited to smaller datasets.

In broad terms, constructing a recogniser involves two stages: detecting region of interest (ROI) and classifying potential signals. The number and complexity of these steps within a pipeline will vary depending on the methodology used. (Lasseck 2014; Sebastián-González et al. 2015; Knight et al. 2020). Region of Interest (ROI) detection identifies potential target signals. One simple and accessible ROI technique is template matching, which can be used as both the ROI identification or classification (Katz, Hafner, and Donovan, 2016). Template matching involves using a measure such as spectral cross-correlation to assess the similarity between one or more reference call patterns and a set of unknown call patterns. Using template matching for classification relies on creating a sufficiently representative call library (Aide et al., 2013; Gibb et al., 2019) and is therefore sensitive to variation in signal structure, i.e., call type and background noise (Brandes, 2008; Katz, Hafner and Donovan, 2016). Combining template matching with machine learning methods reduces the false positive rate of a classifier compared to using template matching alone (Balantic and Donovan, 2020). In this use case, template matching extracts regions of interest (ROIs) that are then classified by an sML algorithm. For example, suppose the template library sufficiently represents intra-specific call type variation with an appropriate cross-correlation score threshold. In that case, template matching will improve the quality of data input into an sML classifier. Several different sML approaches exist; random forest (Brieman, 2001) is one of the most widely used (Tachibana, Oosugi and Okanoya, 2014; Noda, Travieso and Sánchez-Rodríguez, 2016; Raghuram et al., 2016) and performs well at bioacoustic tasks (Weerasena et al., 2018; Ayala-Berdon et al., 2020; Smith-Vidaurre, Araya-Salas, and Wright, 2020).

An essential part of developing a recogniser is selecting the size of data set used to train the sML. There are few concrete guidelines on what constitutes an adequate training dataset size. Sebastián-González et al. (2015) tested the effect of training dataset size on accuracy. They found that a 50% reduction in dataset size resulted in a loss of less than 1% balanced accuracy metric (BAC), suggesting that their chosen sML method (a Support Vector Machine) copes well with small datasets (n=642). However, it is not easy to generalise these results because classifier performance varies on a species-to-species basis. For example, Digby et al. (2013) achieved a recall of 39.8% and precision of 98.1% with 3411 little spotted kiwis (*Apteryx owenii*) calls and 3072 negative cases. In contrast, Sebastián-González et al. (2015) used a maximum of 1285 ‘Amakihi (*Hemignathus viren virens*) and 2785 negative cases and achieved a BAC of 86.5%. Similarly, when attempting to find the best training dataset size to optimise their relative sound level (RSL) method, Knight et al. (2020) found that the common nighthawk (*Chordeiles minor*) had an optimal training dataset size of 10,590 cases. In contrast, the Ovenbird (*Seiurus aurocapilla*) needed only 5,540 cases to achieve similar performance.

A challenge in developing classifiers for PAM studies is the spatiotemporal variation in background noise. For example, various species vocalise concurrently in tropical rainforests in overlapping frequency ranges (Slabbekoorn 2004). The soundscape also varies as biotic factors, such as breeding season, and abiotic factors, such as weather change through time. Plant and animal community composition can also vary spatially across heterogeneous tropical landscapes (Cintra and Naka, 2011; Wardhaugh, Stork, and Edwards, 2014; Ioki *et al*., 2016). These factors make ecosystems like tropical rainforests challenging for PAM (e.g., Heinicke *et al*., 2015). One way to improve classifier performance, which may be especially important when not denoising, is to ensure the training dataset captures the variation in background noise. A sufficiently representative dataset is likely to be very large. Creating such a training dataset is challenging because it requires identifying the many sources of background noise that may be a problem for the classifier. Moreover, determining this kind of background noise would need an *a priori* knowledge that is not available.

## Macaw case study

We present a case study of the development of a PAM recogniser for two sympatric macaw species: the critically endangered great green macaw (GGM - *Ara ambiguus*) (BirdLife International 2020) and the regionally endangered scarlet macaw (SCM - *Ara macao*) (Monge et al. 2016) in northern Costa Rica. Parrots are one of the most endangered families of birds, with 34% of species classified as threatened (IUCN 2020). They are widely distributed globally and native to every continent in the tropics. For many species of conservation concern, there is a lack of data on factors important to conservation planning and policy, such as their distribution and abundance. This limitation is especially concerning in the case of the GGM because they have recently been uplisted to critically endangered (BirdLife International 2020).

Parrots represent a significant challenge for classification tasks due to their wide vocabulary (Taylor and Perrin 2005; Zdenek et al. 2015; Montes-Medina et al. 2016) and context-dependent calls (Bradbury 2003). The two focal species present an additional challenge because their calls are highly similar and difficult to distinguish, even for experts (TL *pers. obvs*). We aimed to construct a GGM-SCM recogniser using PAM data collected from northern Costa Rica during the first six months of 2020. However, due to restrictions arising from the Covid-19 pandemic, we were forced to construct an initial recogniser using only the first six weeks of data. This constraint shaped our workflow and provided a natural experiment that reveals the challenges of creating robust recognisers in acoustically complex environments. Rather than only reporting the final recogniser, we present our work as it developed to provide insights into these challenges and highlight the risks they pose for deployment in the field. The challenges we faced, and lessons learned are directly relevant to other PAM users developing species-specific classifiers in similar settings.

## Materials and Methods

### The audio

#### Survey recordings

We used AudioMoth 1.2 (Hill et al., 2019; LabMaker, Germany) ARUs, which we installed devices on the tallest accessible tree in a 10km grid across the northeast of Costa Rica (Fig 1). They recorded four 30-minute slots throughout the day (7:00-7:30, 10:00-10:30, 13:00-13:30, 16:00-16:30). Recording took place between 27 January 2020 and 30 June 2020. The sampling frequency was not consistent as there was an error in configuring some devices for a portion of the deployment period.

**Figure 1:**
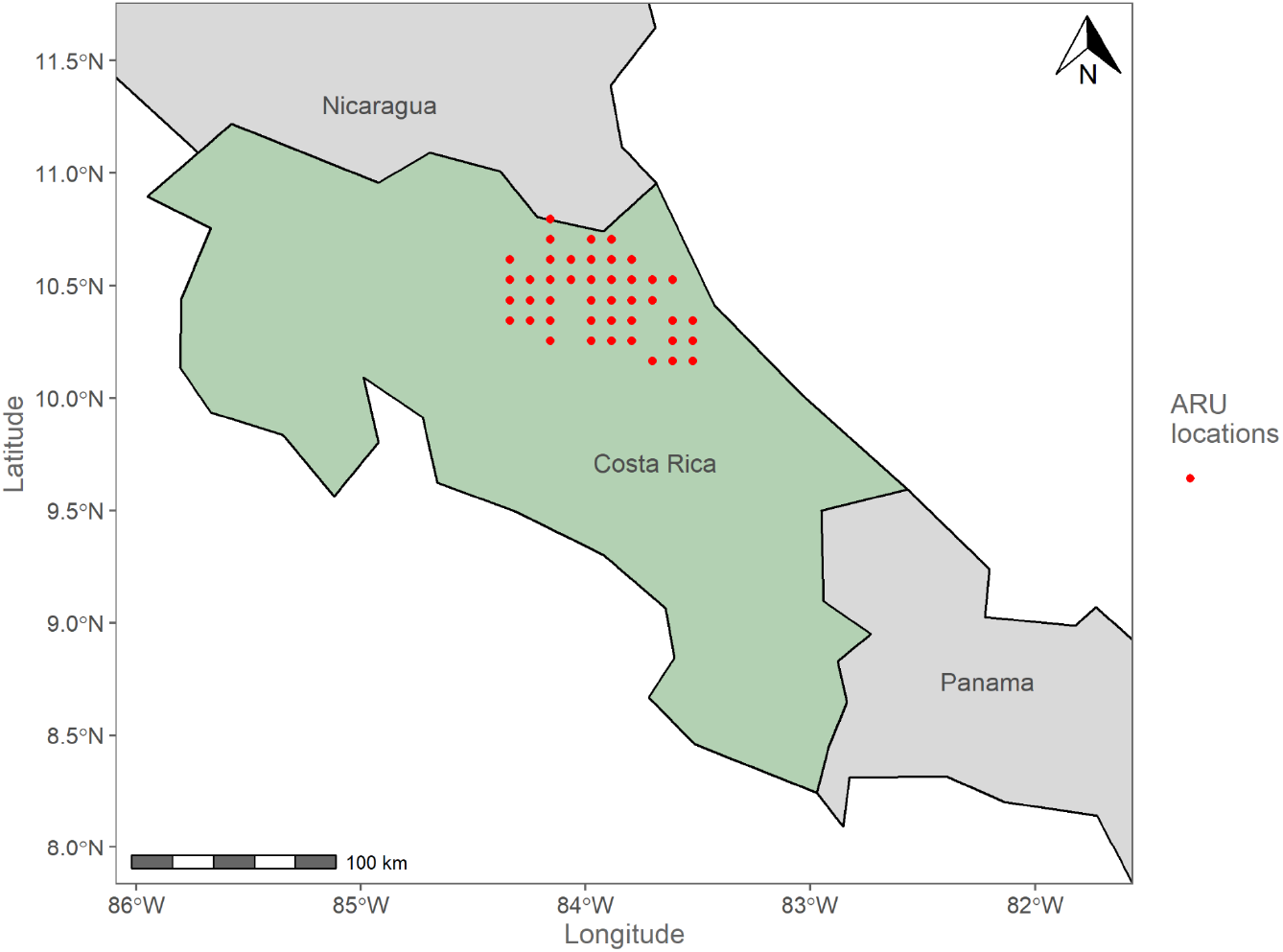
ARU locations in northeastern Costa Rica. ARUs were set out on a 10km grid, holes in the grid are areas that were inaccessible due to dense forest or lack of permission from landowners.

#### High-quality recordings

Template matching requires high-quality recordings of the target species to create reference templates that can be used for spectrogram cross-correlation. We did this with recordings made with a directional microphone Sennheiser ME 67 (Sennheiser electronic GmbH & Co., United Kingdom) and a Roland R-05 Wave/MP3 digital recorder (Roland Corporation, United Kingdom). Recordings were digitised with a 16-bit sampling depth and 48kHz sampling frequency; recordings were saved as Wav files. Recording occurred around known GGM nest sites in northeastern Costa Rica between January and March 2019 (Fig 1). Initially, we selected four recordings containing a total of 157 calls from three different nest sites, as they included single individuals calling to their mate and groups of GGM calling.

#### Acoustic features

We extracted a total of 113 acoustic features using the `warbler` package in R (R Core Team, 2020): 20 measurements of frequency, time, and amplitude parameters, and 93 Mel-frequency cepstral coefficients (MFCCs) (Araya-Salas and Smith-Vidaurre, 2017). MFCCs were initially designed for human speech recognition but have been widely used in bioacoustics (e.g., Loh, Yuan, and Ramli, 2013; Colonna, Gama, and Nakamura, 2016; Noda, Travieso, and Sánchez-Rodríguez, 2016; Salamon *et al*., 2016). MFCCs reduce any signal to a set of coefficients (Colonna, Gama, and Nakamura, 2016) while minimising any loss of biologically relevant information (Davis and Mermelstein, 1980). These are potentially helpful when dealing with large amounts of bioacoustic data.

### Recogniser workflow elements

#### ROI detection: template matching

The high-quality recordings were used to construct templates (n = 157) following Hafner and Katz (2018) (Fig. 2). First, we visually checked templates and removed if they contained any non-target signal (n = 94). With the remaining templates, we evaluated them on a randomly selected recording that included GGM calls. Spectrogram cross-correlation is used to score the similarity of a detection to the template. We set a low default threshold (0.2) to allow flexibility in matching call types and amplitudes. Templates that detected less than 10% (n = 28) or over 200% (n = 17) of the total number of calls were removed. We removed templates with a true-positive to false-positive ratio above 1:5 or if 90% of a template’s detections were the same as another’s. To test their accuracy, we ran the final group of templates (n = 4) over ten randomly selected recordings containing SCM and GGM calls and ten randomly selected recordings without any calls. We set the window length used in the template matching to one second to reflect the maximum call length (1 second).

**Figure 2:**
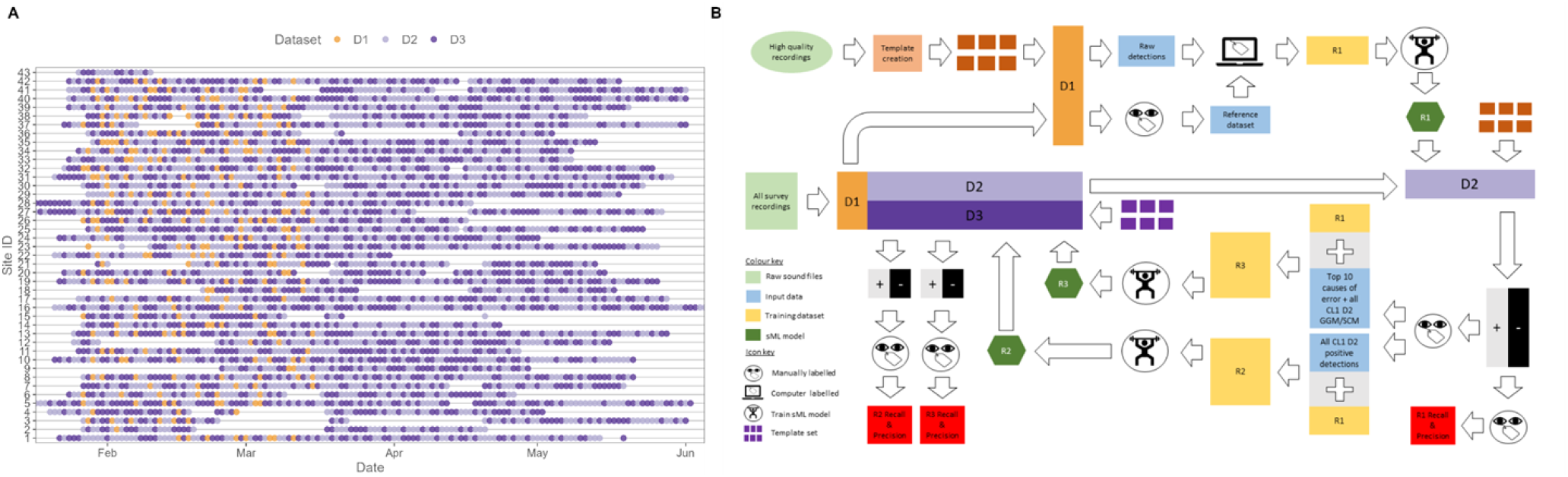
Schematic of classifier development A) represents how data was divided to create the three training/testing datasets. D1 was selected from recordings up to the middle of March when TL had to leave the field due to the pandemic. After April recording devices were left in the field until June and so any failures were not rectified, which resulted in fewer recordings taking place towards the end of the study period. Most sites had issues with recording devices failing at some point; at site 43 the recording device failed and then was stolen so was not re-installed. B) The workflow used to develop each classifier, starting from raw sound files (light green) and resulting in classifiers and performance metrics (red).

#### Signal classification: supervised ML

We used a `tidymodels` workflow (Kuhn and Wickham, 2020) in R (R Core Team, 2020) to train each random forest algorithm. We used a 75:25 split of the training dataset to create the training and test data, meaning that 75% of the training dataset is used to train the classifier, and 25% is withheld to estimate performance. A random forest has three hyperparameters to be tuned: number of trees, number of nodes, and number of variables per node.

### Division of the data

#### Foundation dataset

Due to the global Covid-19 pandemic, data was divided into two; January 27 2020, to March 15 2020, and March 16 2020, to 30 June 2020. From the first set of recordings, we created our first recogniser. We chose an initial target of 1000 calls per species to start the database creation to match the smallest published training dataset size (Sebastián-González *et al*., 2015). First, we manually searched 100 randomly selected recordings to estimate the call rate per 30-minute recording. These were 3.13 GGM and 1.18 SCM calls per recording. Thus, our foundation dataset was a total of 848 recordings. We created the dataset by sampling recordings across all sites and survey times using Sobol sequences to create a pseudo-random sample (Sobol, 1967; Antonov and Saleev, 1979) (Fig. 2). We manually labelled these to generate the reference dataset; 2773 GGM calls and 843 SCM calls. For manual labelling, we loaded recordings into Raven Lite (Bioacoustics Research Program, 2016) and visually examined their spectrograms for putative target species calls. Suspected target calls were confirmed by listening to them. We ran our template set over the 848 recordings and used the reference dataset to label all positive cases; the remaining detections were labelled negative.

#### Build & Validation datasets

When we received the second set of recordings from the field (March 16 2020, to 30 June 2020), we combined them with the recordings not used in the foundation dataset. We then divided these recordings into two sets, one that we used to evaluate the accuracy of the recogniser one and build a subsequent recogniser. We could use the second use to assess the accuracy of any other recognisers. Finally, to ensure an even spatial and temporal sampling of the recordings in each set, we again used Sobol sequences to create a pseudo-random sample (Sobol, 1967; Antonov and Saleev, 1979). We ended up with two equally sized datasets (build: n = 8084 / validation: n = 8084 - Fig. 2).

### Recogniser one

To create the training dataset for recogniser one, we used all positive cases in our foundation dataset plus an equal number of negative cases. The negative cases were selected by pseudo-random sampling across sites and survey periods using Sobol sequences (Sobol, 1967; Antonov and Saleev, 1979). Therefore recogniser one’s training dataset consisted of 4247 GGM calls, 1393 SCM calls, and 5710 negative cases. The number of positive cases was higher than the number of manually labelled reference data because overlapping calls were only recorded once when manually labelling them. However, we designed the template matching detections not to overlap because the multiple templates could detect the same call. Therefore these continuous calls would be detected as multiple cases rather than a single manually labelled case. We used this dataset to train recogniser one.

### Recogniser performance

We used recall and precision as our performance metrics. Recall is a measure of the false negative (FN) rate (Recall = FN / TP + TN) and precision is a measure of false positive (FP) rate (Precision = FP / FP + TP). We initially used the performance metrics estimated on the 25% withheld test data to assess the accuracy of the random forest. We changed this approach after applying recogniser one to the build dataset; we found that performance was significantly worse than the initial estimates suggested. We decided to manually check all positive cases and conduct a power analysis to determine how many negative cases we needed to check manually. For this, we needed the false-negative effect size:

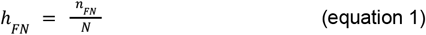

Where *n_FN_* is the number of false-negative cases, and *N* is the total number of all cases, as well as significance level (0.05) and power (0.95). As we used the initial test dataset to calculate the effect sizes, a large number of negative cases needed to be checked to be able to detect such small effect sizes. We then used false-positive, false-negative, true-positive and true-negatives from these two manually checked datasets to evaluate actual recogniser performance.

### Sources of error

Our initial assumption was that the macaws’ calls are significantly distinct, so the recognisers would not have an issue distinguishing these calls from other sounds. When applied to the build dataset, the poor performance of recogniser one suggested this was not the case. So, we went back through all previously checked positive detections and subsequently, while checking more false-positives, where possible, we labelled each case to species or genera level where possible. Finally, we investigated how sources of error change over time and how this affects recogniser precision.

### Recognisers two & three

We used all of the manually checked positive cases used to evaluate recogniser one and the dataset used to train recogniser one to train recogniser two. We also trained the third recogniser on a subsection of the whole large training dataset, using 1000 cases of the top 7 sources of error. Finally, we completed the negatives by randomly selecting them from other sources of error, so the total negative cases were equal to the total GGM. We did this because we wanted to know if being specific about the origins of non-target signals would be a way to increase accuracy while limiting the size and, therefore, the time needed to train the recogniser. After this, we applied the two newly trained recognisers to the validation dataset and estimated performance.

## Results

### ROI detection: template matching

Across the foundation, build, and validation datasets, the template set made 2072080 detections, a mean of 121.75 detections per recording (n = 17020). When we look individually at the foundation dataset, the mean detection rate was 119.98 per recording (n = 848). The mean number of GGM calls found when we manually labelled the foundation dataset was 3.27 (n = 848), suggesting that the template matching step captured a lot of non-target signals.

### Recogniser one

We were encouraged when the performance of recogniser one, estimated using the 25% withheld test data, was high in both recall (GGM = 0.92 / SCM = 0.85) and precision (GGM = 0.93 / SCM = 0.96 - Fig. 3). When applied to the build dataset, recogniser one made 37639 positive detections (35814 GGM and 1825 SCM), 4161 were true GGM and 201 true SCM when we manually checked them. A power analysis using the SCM false negative effect size, as it was the lowest of either target species (GGM = 0.0327, SCM = 0.0177), determined that we needed to manually check a minimum of 41514 negative cases to detect this effect size. Finally, we reviewed all positive cases (n = 37639) to estimate the performance of the recogniser one when exposed to the full diversity of ambient noise across sites and seasons. When applied to this build dataset, classifier precision degraded (GGM = 0.12 / SCM = 0.11) significantly for both species. Recall declined for the SCM (0.04), whereas it increased for the GGM (0.98 - Fig. 3).

**Figure 3:**
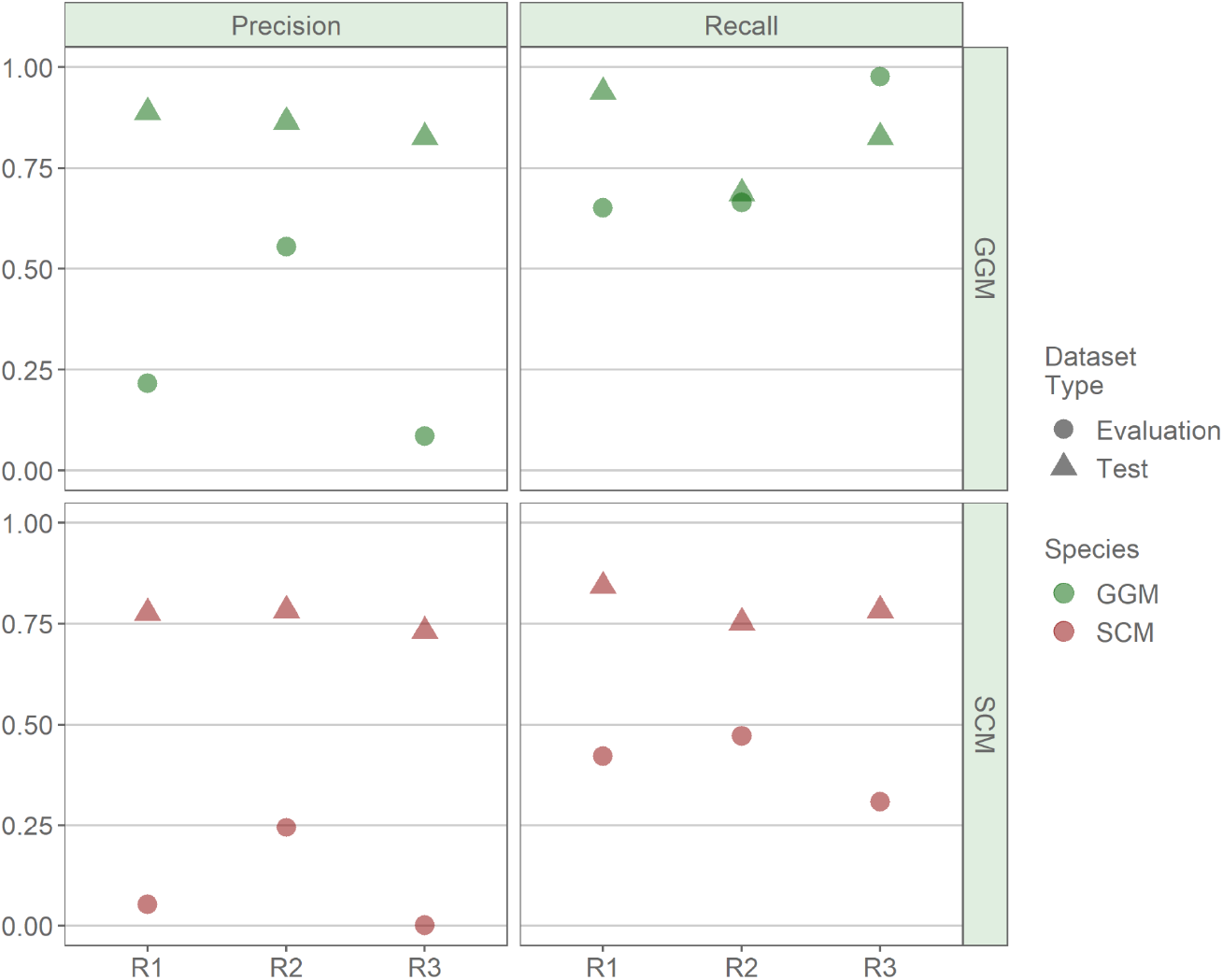
Classifier performance metrics show the improvement between CL1 and CL3. There is a trade-off between precision and recall, when precision increases, recall decreases and *visa-versa*. There is a constant large difference between evaluation and validation precision metrics, this is the smallest for CL3 but demonstrates the potential pitfalls of using default validation datasets to estimate the performance of classifiers.

### Recogniser two

We combined the 4362 (4161 GGM + 201 SCM) detections made by recogniser one in the build dataset with the 5640 (4247 GGM + 1393 SCM) detections from the foundation dataset. Recogniser two was trained with these (8408 GGM + 2112 SCM) and all non-target positive detections from recogniser one (n = 31999) plus the negative cases used to train recogniser one (n = 5710). Initial performance was estimated using the test data. Both recall (GGM = 0.69 / SCM = 0.75) and precision (GGM = 0.86 / SCM = 0.78) was lower than the performance of recogniser one (Fig. 3). This is likely due to the size and spatio-temporal scale of the training dataset being larger and capturing more background noise, therefore being more representative. Hence, performance estimates are closer to reality. We manually checked all positive detections (n = 6390). Using the SCM false negative effect size as it was the smallest (GGM = 0.0498, SCM = 0.0104), a power analysis determined we needed to manually check a minimum of 119759 negative cases to detect this effect size. Recall for the GGM did not change significantly (0.66), whereas precision did drop (0.56), but it was not by as much as for recogniser one. SCM performance dropped for both metrics (Recall = 0.66 / Precision = 0.24), but the decline was less severe than for recogniser one. The discrepancy between the two species’ performance is due to the large difference in the training dataset size for each species.

### Sources of error

The sources of error varied through time (Fig. 4) and space (Fig. 5). However, we can see a noticeable change in the dominant source of error after the period used to create the recogniser one (Fig. 4B). This suggests that, even though performance is poor, the temporal variation in background noise is driving the even worse performance later in the season (Fig. 4A). Unidentified passerines stay a constant proportion of the error, whereas Amazona *spp*. decrease after this point, whereas clay-coloured thrush (*Turdus grayii*) and chickens (*Gallus domesticus*) increase. This is probably because the breeding season of the clay-coloured thrush begins after March, whereas the breeding season for Amazona *spp*. ends around this time.

**Figure 4:**
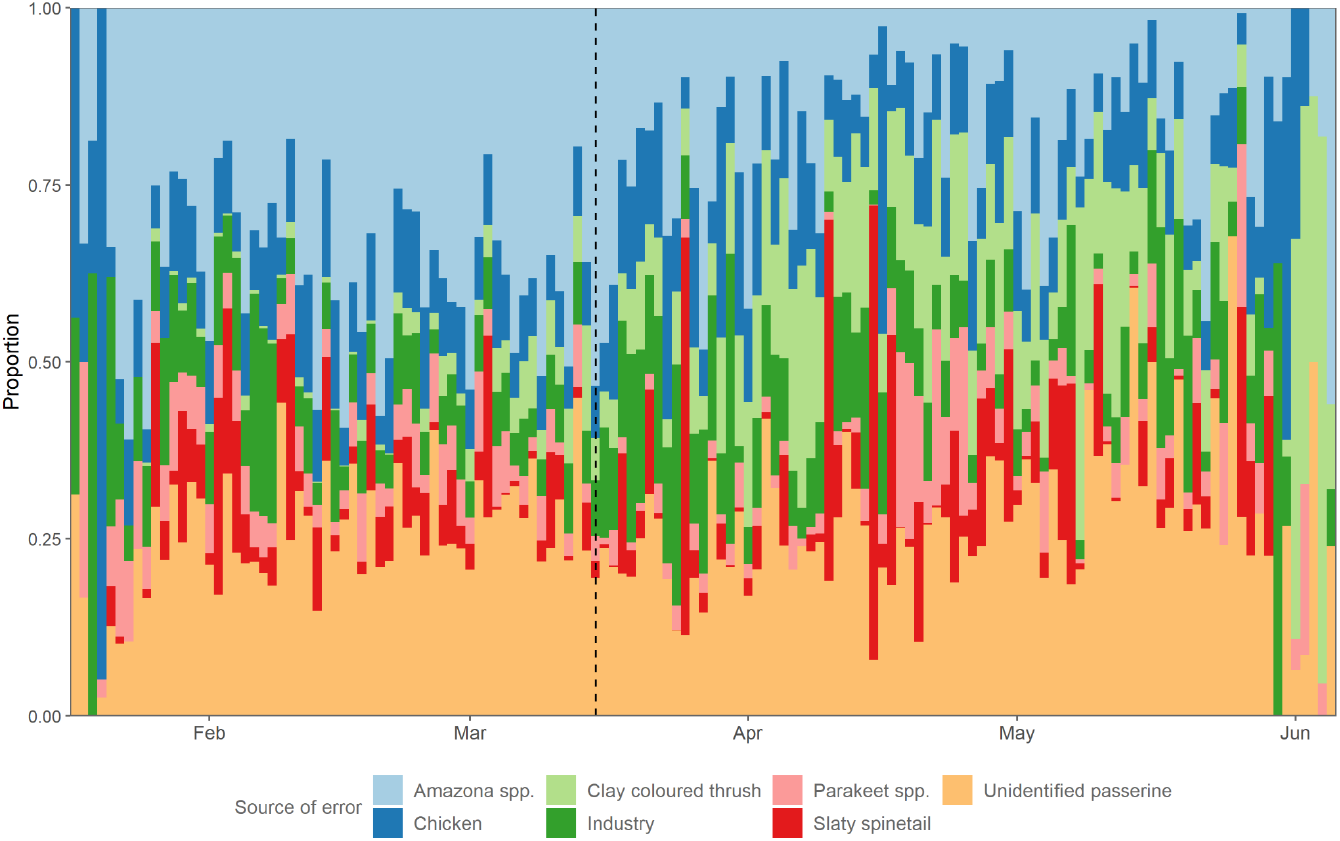
The temporal change in source background noise in the false positive detection of recogniser one in the evaluation dataset across the study period. There is a distinct change in the proportions of *Amazona* spp., clay colour thrush and chicken after the time period that the recogniser one’s training dataset was selected (dashed line).

**Figure 5:**
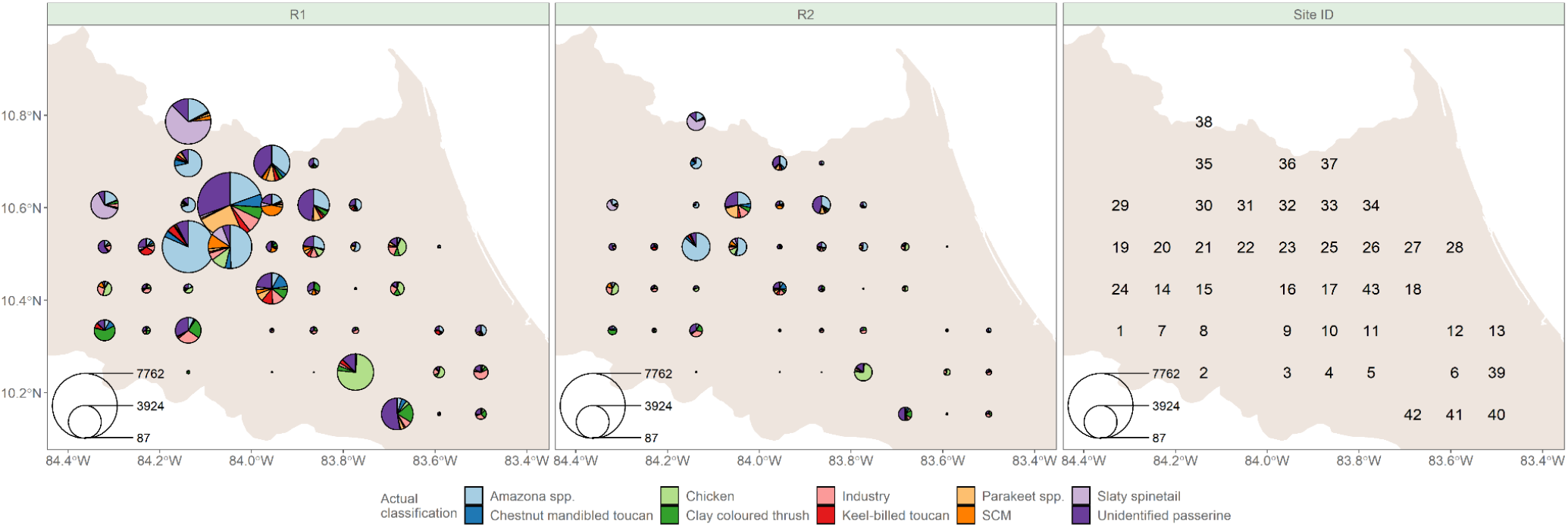
Site-level variation in the overall top 10 sources of error, SCM and GGM detections from recogniser one (R1) and recogniser two (R2). There is a clear difference in the sources of error across the study sites, with northern sites being dominated by *Amazona* spp., unidentified passerines and at sites 29 and 38, the slaty spinetail (*Synallaxis brachyura*). Overall false positives were reduced by 90% from R1 to R2 and at each site, the variation in sources of error was also generally reduced.

Overall, false-positive detections from all sources were reduced by recogniser two compared to recogniser one, apart from the misclassification of GGM as SCM (Fig. 5). Recogniser one’s false-positive detections at many sites are dominated by one or two sources of error (e.g. site 5, 1 and 29 - Fig. 5). In contrast, there is a smaller number with a wide variety of sources of error (e.g. site 16, 20 and 25). However, there is a distinct spatial variation in the types of sources of error. For example, misclassifications are caused mainly by other wild bird species in the northern sites, whereas there is more anthropogenic noise (chainsaw, industry) in the south and east. Although a wild bird, the clay colour thrush is often strongly associated with areas of human disturbance (Dyrcz 1983), and our results support this association. Spatial variation in the source of error demonstrates that capturing this is critical in creating a representative training dataset.

### Recogniser three

We constructed the training dataset for recogniser three using Amazona *spp*., unidentified passerine, chicken, industry, clay-coloured thrush, slaty spinetail and parakeet *spp*.. One thousand cases of each were combined with 1408 randomly selected other negative cases to make a negative dataset equal to that of the GGM. The final training dataset for recogniser three was 8408 GGM, 2112 SCM and 8408 negatives. When applied to the validation dataset the recogniser made 26480 positive detections (GGM = 12536 / SCM = 13944). Both recall (GGM = 0.83 / SCM = 0.78) and precision (GGM = 0.83 / SCM = 0.73) were high when estimated using the validation dataset. We checked all positive cases and performed a power analysis using the SCM, the smallest false negative effect size again (GGM = 0.0774 / SCM = 0.0241). This determined that a minimum of 22374 negative cases had to be checked. Performance dropped significantly for precision (GGM = 0.09) / SCM = 0.001), while recall increased for the GGM (0.97) but decreased for the SCM (0.31).

## Discussion

We have demonstrated that users must be careful when preparing a training dataset for a PAM recogniser. A training dataset that captures relevant background noise variation across space and time is required to construct a robust recogniser. Our workflow addresses this by using an initial small recogniser to create a training dataset that captures non-target signals driving false-positive detections. This method is similar to the “basic recognition model” approach used by Buxton and Jones (2012), whereby a small amount of data is used to train a basic model, which is then used to search for more positive training data in other recordings. The main difference is that we extended this approach to gather positive and negative cases.

We initially assumed GGM and SCM calls were sufficiently distinctive that using recordings from a narrow window in the breeding season would allow us to construct a robust recogniser. However, the performance of our initial breeding season recogniser was very poor when applied to data from outside the breeding season due to the increased diversity of the soundscape. Importantly, this variation is difficult to capture without using a large, manually labelled dataset. This would have required us to label over 9000 30-minute recordings. Incorporating the positive detections and previous training datasets to train a second classifier significantly increases precision whilst losing only a small level of recall performance. This method is similar to the workflow of Balantic and Donovan (2020), using template matching and machine learning together to reduce false positives. The main benefit of our approach is the relatively simple pipeline structure. Nonetheless, we found that iterative labelling and training were required to create a useable recogniser when the target species exhibits considerable call type variation and high spatiotemporal variation in background noise.

Our enhanced recogniser that included seasonal variation (R2) still performed poorly compared to many other machine learning PAM studies (Jahn et al. 2017; Bravo et al. 2017; Knight and Bayne 2019; Knight et al. 2020; Balantic and Donovan 2020; Gillings and Scott 2021). However, our results are comparable to similar studies in tropical environments (Swiston and Mennill 2009; Heinicke et al. 2015). This reduced performance likely reflects the highly diverse soundscape arising from the high diversity of species with overlapping signal frequencies (Slabbekoorn 2004) and anthropogenic noise (Slabbekoorn and Peet 2003) in tropical settings. Both factors were apparent in our study region in the northeast of Costa Rica, which encompasses primarily cropland and urban in the south, and mainly cattle pasture and forest land use types in the north (Fagan et al. 2013; Jadin et al. 2016; Karra et al. 2021).

Another important finding was that the performance of the breeding season recogniser was hugely estimated. Practitioners should consider this when creating classifiers for large-scale rollout. Here we have been explicit in describing the data used to assess model performance, but this is not always the case (Heinicke et al., 2015). It is unclear how many published recognisers would perform in a real-world PAM study, especially if studies use high-quality recordings to create their classifier (Bardeli et al. 2010; Buxton and Jones 2012; Priyadarshani et al. 2018). Manually checking positive detections and using power analysis to determine how many negative cases need to be reviewed is a more labour-intensive task but gives a more accurate assessment of real-world performance.

We did not succeed in creating a recogniser that can be used without manually checking outputs. This is not uncommon. Even when classifiers have high-performance metrics, manually checking positive classifications to filter out false positives is often done before downstream analyses of species abundance and distribution (Buxton and Jones, 2012; Zwart et al., 2014; Colbert et al., 2015; Kalan et al., 2015; Sidie-Slettedahl et al., 2015). A way to deal with this need for manual checking is to use statistical methods that can reduce the amount of data needed for validation (Knight et al., 2020), account for false positives (Banner et al., 2018), false negatives (MacKenzie et al., 2002) or both (Chambert et al., 2018; Wright et al., 2020). Our method reduced the number of false positives that must be manually validated by 80%, from 3.95 per recording (n = 33243) in R1 to 0.75 per recording (n = 6309) in R2. Therefore, although it is still necessary to manually check all positive detections when the final recogniser runs on new data, this represents a significant reduction in the effort needed to clean the data.

Denoising can increase classifier accuracy (Stowell et al. 2016). Interestingly, denoising is not often used, even when studies report the high performance of recognisers. This supports the argument made by Priyadarshani, Marsland and Castro (2018) that many published PAM studies are done in low-noise environments, using species with simple calls. The primary barrier to denoising is that few simple, user-friendly techniques are available, so many classifiers do not use them (e.g. Sebastián-González *et al*., 2015; Balantic and Donovan, 2020; Knight *et al*., 2020). We provide an alternative way to tackle the issue of background noise without using denoising. However, it does not entirely deal with the problem as our recogniser performance is not comparable to the best-published recognisers.

## Future work

We could improve our classifier in several ways. Effective template matching requires a sufficiently representative call library (Aide et al., 2013; Gibb et al., 2019). We set the matching threshold very low to help our classifier deal with intra- and inter-call type variation of the GGM and SCM. Although we did not miss any target signal, the random forest classified a large volume of data. Investing extra time in developing and refining the template set would likely reduce the time needed to review calls and improve performance by reducing the size of the initial dataset.

Training dataset imbalance is common in machine learning, not just in bioacoustics (Salamon and Bello 2017). Methods such as data augmentation can provide an excellent solution to this problem (Stowell et al., 2019). Our training dataset was very imbalanced, and the quantity of SCM training data was only ~25% of the GGM training dataset. This resulted in the performance of SCM being significantly lower than that of the GGM. Our current workflow uses a standard window size of one second, within which the call may occupy only a fraction. Therefore, we would have to change how we structure our pipeline to accommodate data augmentation, as many data augmentation techniques need to have the start and end of the target signal (Salamon and Bello 2017). Being able to automatically find the beginning and end of a target signal would be a massive step toward facilitating data augmentation. It would also help reduce the effect of background noise on random forest accuracy by reducing the amount of background noise present in each window to be classified.

## Conclusions

Passive acoustic monitoring has great potential to enable the scaling up of biodiversity monitoring to inform policy and conservation strategies. There is an increasing need for simple methods to automate or semi-automate data extraction from PAM surveys (Marques et al., 2013; Stowell et al., 2016). The development of generalised deep-learning classifiers will make PAM a simple and accessible tool for biodiversity monitoring. However, their development may still be a long way off, and species-specific recognisers can be more accurate (Priyadarshani et al., 2018). Custom recognisers also allow practitioners to tailor them to their specific needs. Developing such recognisers is challenging, and although there is an increasing amount of literature on different methods to create classifiers, these are often small-scale studies in temperate environments (Sugai *et al*. 2019) that rely on high-quality recordings of species with simple calls (Priyadarshani et al. 2018). We have demonstrated a simple workflow that can provide users with an accessible but time-consuming way to create a PAM recogniser for acoustically diverse environments.

## Bibliography

Balantic CM, Donovan TM. 2020. Statistical learning mitigation of false positives from template-detected data in automated acoustic wildlife monitoring. Bioacoustics. 29(3):296–321. https://doi.org/10.1080/09524622.2019.1605309

Bardeli R, Wolff D, Kurth F, Koch M, Tauchert K-H, Frommolt K-H. 2010. Detecting bird sounds in a complex acoustic environment and application to bioacoustic monitoring. Pattern Recognit Lett. 31(12):1524–1534. https://doi.org/10.1016/j.patrec.2009.09.014

Bravo CJC, Berríos RÁ, Aide TM. 2017. Species-specific audio detection: a comparison of three template-based detection algorithms using random forests. PeerJ Comput Sci. 3:e113. https://doi.org/10.7717/peerj-cs.113

Buxton RT, Jones IL. 2012. Measuring nocturnal seabird activity and status using acoustic recording devices: applications for island restoration. J Field Ornithol. 83(1):47–60. https://doi.org/10.1111/j.1557-9263.2011.00355.x

Cintra R, Naka LN. 2011. Spatial Variation in Bird Community Composition in Relation to Topographic Gradient and Forest Heterogeneity in a Central Amazonian Rainforest. Int J Ecol. 2012:e435671. https://doi.org/10.1155/2012/435671

Dyrcz A. 1983. Breeding ecology of the Clay-coloured Robin Turdus grayi in lowland Panama. Ibis. 125(3):287–304. https://doi.org/10.1111/j.1474-919X.1983.tb03115.x

Fagan ME, DeFries RS, Sesnie SE, Arroyo JP, Walker W, Soto C, Chazdon RL, Sanchun A. 2013. Land cover dynamics following a deforestation ban in northern Costa Rica. Environ Res Lett. 8(3):034017. https://doi.org/10.1088/1748-9326/8/3/034017

Gillings S, Scott C. 2021. Nocturnal flight calling behaviour of thrushes in relation to artificial light at night. Ibis [Internet]. [accessed 2021 May 5]. https://doi.org/10.1111/ibi.12955

Heinicke S, Kalan AK, Wagner OJJ, Mundry R, Lukashevich H, Kühl HS. 2015. Assessing the performance of a semi-automated acoustic monitoring system for primates. Methods Ecol Evol. 6(7):753–763. https://doi.org/10.1111/2041-210X.12384

Ioki K, Tsuyuki S, Hirata Y, Phua M-H, Wong WVC, Ling Z-Y, Johari SA, Korom A, James D, Saito H, Takao G. 2016. Evaluation of the similarity in tree community composition in a tropical rainforest using airborne LiDAR data. Remote Sens Environ. 173:304–313. https://doi.org/10.1016/j.rse.2015.07.024

Jadin I, Meyfroidt P, Lambin EF. 2016. International trade, and land use intensification and spatial reorganization explain Costa Rica’s forest transition. Environ Res Lett. 11(3):035005. https://doi.org/10.1088/1748-9326/11/3/035005

Jahn O, Ganchev TD, Marques MI, Schuchmann K-L. 2017. Automated Sound Recognition Provides Insights into the Behavioral Ecology of a Tropical Bird. PLOS ONE. 12(1):e0169041. https://doi.org/10.1371/journal.pone.0169041

Karra K, Kontgis C, Statman-Weil Z, Mazzariello JC, Mathis M, Brumby SP. 2021. Global land use / land cover with Sentinel 2 and deep learning. In: 2021 IEEE Int Geosci Remote Sens Symp IGARSS. [place unknown]; p. 4704–4707. https://doi.org/10.1109/IGARSS47720.2021.9553499

Katz J, Hafner SD, Donovan T. 2016. Tools for automated acoustic monitoring within the R package monitoR. Bioacoustics. 25(2):197–210. https://doi.org/10.1080/09524622.2016.1138415

Knight EC, Bayne EM. 2019. Classification threshold and training data affect the quality and utility of focal species data processed with automated audio-recognition software. Bioacoustics. 28(6):539–554. https://doi.org/10.1080/09524622.2018.1503971

Knight EC, Sòlymos P, Scott C, Bayne EM. 2020. Validation prediction: a flexible protocol to increase efficiency of automated acoustic processing for wildlife research. Ecol Appl. 30(7):e02140. https://doi.org/10.1002/eap.2140

Lasseck M. 2014. Large-scale identification of birds in audio recordings.:11.

Priyadarshani N, Marsland S, Castro I. 2018. Automated birdsong recognition in complex acoustic environments: a review. J Avian Biol. 49(5):jav-01447. https://doi.org/10.1111/jav.01447

Salamon J, Bello JP. 2017. Deep Convolutional Neural Networks and Data Augmentation for Environmental Sound Classification. IEEE Signal Process Lett. 24(3):279–283. https://doi.org/10.1109/LSP.2017.2657381

Sebastián-González E, Pang-Ching J, Barbosa JM, Hart P. 2015. Bioacoustics for species management: two case studies with a Hawaiian forest bird. Ecol Evol. 5(20):4696–4705. https://doi.org/10.1002/ece3.1743

Slabbekoorn H. 2004. Habitat-dependent ambient noise: Consistent spectral profiles in two African forest types. J Acoust Soc Am. 116(6):3727–3733. https://doi.org/10.1121/1.1811121

Slabbekoorn H, Peet M. 2003. Birds sing at a higher pitch in urban noise. Nature. 424(6946):267–267. https://doi.org/10.1038/424267a

Stowell D, Petrusková T, Šálek M, Linhart P. 2019. Automatic acoustic identification of individuals in multiple species: improving identification across recording conditions. J R Soc Interface. 16(153):20180940. https://doi.org/10.1098/rsif.2018.0940

Stowell D, Wood M, Stylianou Y, Glotin H. 2016. Bird detection in audio: A survey and a challenge. In: 2016 IEEE 26th Int Workshop Mach Learn Signal Process MLSP. p. 1–6. https://doi.org/10.1109/MLSP.2016.7738875

Swiston KA, Mennill DJ. 2009. Comparison of manual and automated methods for identifying target sounds in audio recordings of Pileated, Pale-billed, and putative Ivory-billed woodpeckers. J Field Ornithol. 80(1):42–50. https://doi.org/10.1111/j.1557-9263.2009.00204.x

Wardhaugh CW, Stork NE, Edwards W. 2014. Canopy invertebrate community composition on rainforest trees: Different microhabitats support very different invertebrate communities. Austral Ecol. 39(4):367–377. https://doi.org/10.1111/aec.12085

White E. 2018. Minimum Time Required to Detect Population Trends: The Need for Long-Term Monitoring Programs. BioScience. 69. https://doi.org/10.1093/biosci/biy144

