## Supplementary Materials 1 for "Inadequate sampling of the soundscape leads to overoptimistic estimates of recogniser performance: a case study of two sympatric macaw species"

PAM Recogniser training


### PAM Recogniser training

#### TL

#### 2022-11-21

The code included here will train a random forest classifier,
calculate performance and create figure X. This classifier is one part
of the recogniser created for Lewis et al. 2023.

The call templates that created the initial training dataset and that
was the input to the first classifier can be found in the associated
files.

### Packages

```
X <- c('tidymodels','workflows','tune','ranger','rsample', 'parallel', 'themis',
       'tidyverse','randtoolbox','pROC','pwr','ggpubr','doParallel','Rraven')
invisible(lapply(X, library, character.only = TRUE))
```

### ML Training

#### Load data

This data already has the features extracted. To get these features
`warbleR` functions `specan` and `mfcc`
were used.

```
# template matched csv
data <- read.csv("./Data/Training_data_features.csv") %>%
  mutate(Common = ifelse(Common == "GGM", "GGM", 
                      ifelse(Common == "SCM", "SCM", "no"))) 

final_df <- data %>% 
  select(5:ncol(.))

print(final_df %>% count(Common))
```

```
##   Common     n
## 1    GGM  8408
## 2     no 31017
## 3    SCM  2112
```

#### Train

```
# get usable cores
cores <- detectCores() - 1

# inital split
data_split <- initial_split(final_df, prop = 3/4)

# split 75/25
data_train <- training(data_split)
data_test <- testing(data_split)
data_xv <- mc_cv(data_train)

# defining a recipe to allow specification of variable roles etc and pre-processing
data_recipe <- recipe(Common ~ ., data = data_train) %>%
  step_normalize(all_numeric()) %>%
  step_smote(Common)

# ---- Random Forest ----
# now we specify the rf model we will be using
rf_model <-   # specify that the model is a random forest
  rand_forest() %>%
  set_args(mtry = tune(), # specify that the `mtry` parameter needs to be tuned
           min_n = tune(),
           trees = 1000) %>% # this is not set as we will be tuning later
  set_engine("ranger", importance = "impurity") %>% # select the engine/package that underlies the model
  set_mode("classification") # choose either the continuous regression or binary classification mode

# now put all together in a workflow
rf_workflow <- workflow() %>%
  add_recipe(data_recipe) %>% # add the recipe
  add_model(rf_model) # add the model

# define the tuning grid
rf_grid <- grid_regular(
  mtry(range = c(10, 30)),
  min_n(range = c(2, 10)),
  levels = 5)

doParallel::registerDoParallel(cores = cores)

# extract results
set.seed(345)
rf_tune_results <- rf_workflow %>%
  tune_grid(resamples = data_xv, #CV object
            grid = rf_grid, # grid of values to try
            metrics = metric_set(accuracy, roc_auc)) # metrics we care about

# finalise the workflow
# extract the best value for the tune parameters
rf_final <- rf_tune_results %>%
  select_best(metric = "roc_auc")

# Then we can add this parameter to the workflow
rf_final_workflow <- rf_workflow %>%
  finalize_workflow(rf_final)

# finalising the model
rf_model <- fit(rf_final_workflow, final_df) 

saveRDS(rf_model, "./Data/rf_model.rds")
print("OUTPUTS SAVED as ./Data/rf_model.rds", "\n")

# evaluate the model on the test dataset
rf_fit <- rf_final_workflow %>%
  last_fit(data_split) # fit on the training set and evaluate on test set
```

#### Model outputs

We are only interested in the model predictions as the default
performance metrics aren’t done by class, only as a whole. So, we will
calculate performance using the function below.

```
rf_preds <- rf_fit %>% 
  collect_predictions() %>% 
  bind_rows() 

write_csv(rf_preds, "./Data/R2_RF_evaluation.csv")
```

To accurately calculate performance metrics I have a function:

```
## Metrics function -----
metric_fun <- function(data){
  
  # create list for each target species
  y <- rep(list(list()), nrow(as.data.frame((unique(data[,6])))))
  
  for (i in seq_along(y)) {
    # assign 1 or 0 for the actual cases state
    x <- within(data, actual <- ifelse(data[,5] == paste0(as.data.frame(unique(data[,6]))[i,]), 1, 0))
    
    # assign 1 or 0 for the predicted case state
    x <- within(x, pred <- ifelse(data[,6] == paste0(as.data.frame(unique(data[,6]))[i,]), 1, 0))
    
    # assign classification state
    y[[i]] <- within(x, correct <- ifelse(actual == 1 & pred == 1, "TP",
                                          ifelse(actual == 1 & pred == 0, "FN",
                                                 ifelse(actual == 0 & pred == 1, "FP", "TN"))))
    
    # add the target species into the dataframe
    y[[i]]$target <- paste0(as.data.frame(unique(data[,6]))[i,])
  }
  
  y <- bind_rows(y)
  
  y %>% count(target, correct)
}
```

```
rf_preds <- read_csv("./Data/R2_RF_evaluation.csv")

rf_metrics <- rf_preds %>% 
  select(-id) %>% 
  metric_fun(.) %>% 
  pivot_wider(names_from = correct, values_from = n) %>% 
  mutate(Precision = TP / (TP + FP),
         Recall = TP / (TP + FN))
```

### Use model on larger dataset

Here we need to run the trained model on a larger dataset get more
accurate performance metrics. To do that we need to load in another
dataset. After this we will have to manual check the detections in
RavenLite.

#### Predicting

```
# load the model
rf_model <- readRDS("./Data/R2.rds")

# predict using this dataset
pred_data <- read_csv("./Data/D3_detections.csv")

predictions <- predict(rf_model, pred_data %>% select(9:ncol(.)), type = "class") %>% 
  bind_cols(read_csv("./Data/D3_detections.csv")  %>% 
              select(1,5:7), .) %>% 
  mutate(start = round(start, 2),
         end = round(end, 2))
```

We will now export all the positive detections for manual checking
and labelling. This section will not be possible without all the
associated recordings, however the outputs of this code chunk (selection
tables) are included in the project folder.

```
predictions %>% filter(.pred_class != "no") %>% 
  arrange(sound.files, start) %>% # must do this or the function does not export everything in the correct selection table and mixes things up
  exp_raven(., 
            path = "./Data/Selection_tables/Check_pos/",
            sound.file.path = "Z:/childs_group1/ggm_audio/3.survey_files/", 
            single.file = F, 
            parallel = T)
```

##### Power analysis

From our performance metrics calculated above we can see there are
different effect sizes

```
# calculate effect sizes
effect_all <- rf_metrics %>% 
  select(1:5) %>% 
  mutate(total = FP + FN + TP + TN,
         FN_effect = FN / total,
         FP_effect = FP / total)

# this will give the number of cases that need to be checked to be 
# able to detect the SCM (smallest) effects
# false negative
FN_power <- pwr.p.test(h = effect_all$FN_effect[3], sig.level = 0.05, power = 0.95)
FN_detects <- FN_power$n

# now we need to work out how many recordings that is, so we need
# to work out the mean detections (positive and negative) per recordings

# negative
negatives <- predictions %>% 
  filter(.pred_class == "no") %>% 
  summarise(detections = nrow(.),
            sound.files = length(unique(sound.files)),
            mean = detections / sound.files)

n.sample <- ceiling(FN_detects / negatives$mean)

# now create df of all negative sound.files 
n.sound.files <- predictions %>% 
  filter(.pred_class == "no") %>% 
  distinct(sound.files)

# dataframe of just the negative cases
data_neg <- predictions %>% 
  filter(.pred_class == "no")

# randomly sample
prs <- randtoolbox::sobol(n = n.sample, dim = 1, seed = 4711)
  
# turn these real numbers into integer indices
prs <- ceiling(prs * nrow(n.sound.files)) 
  
# select just the prs row numbers
n_final <- n.sound.files[prs,]
```

Select all the selection tables in these sound.files

```
data_neg %>% 
  filter(sound.files %in% n_final$sound.files) %>% 
  group_by(sound.files) %>% 
  mutate(selec = row_number()) %>% 
  select(sound.files, start, end, selec) %>% 
  ungroup() %>% 
  arrange(sound.files, start) %>% 
  exp_raven(., 
            path = "./Data/Selection_tables/Check_neg/",
            sound.file.path = "Z:/childs_group1/ggm_audio/3.survey_files/", 
            single.file = F, 
            parallel = T)
```

Now we have to load these sound files into RavenLite and manually
check and label them.

#### Performance

As we can’t manually label the calls I have included the metrics
output as a .csv which we can load in and make the graphs

```
metrics <- rf_metrics %>% 
  select(target, Precision, Recall) %>% 
  pivot_longer(cols = 2:3, names_to = "Metric", values_to = "values") %>% 
  mutate(Type = "Evaluation",
         Recogniser = "R2") %>% 
  bind_rows(., read_csv("./Data/metrics.csv")) %>% 
  filter(target != "no")
```

```
ggplot(metrics) +
  geom_point(aes(x = Recogniser, y = values, colour = target, shape = Type), size = 4, alpha = 0.5) +
  facet_grid(target~Metric) +
  scale_colour_manual(values = c("darkgreen", "darkred")) +
  scale_y_continuous(limits = c(0,1)) +
  theme(panel.grid.major.x = element_blank(),
        panel.grid.minor.y = element_blank(),
        panel.grid.major.y = element_line(colour = "gray80"),
        panel.background = element_blank(),
        panel.border = element_rect(fill = NA, colour = "gray40"),
        axis.ticks = element_line(colour = "gray40"),
        axis.title.y = element_blank(),
        axis.title.x = element_blank(),
        axis.text = element_text(size = 12, colour = "gray40"),
        axis.text.x = element_text(size = 12, colour = "gray40"),
        legend.title = element_text(size = 12, colour = "gray40"),
        legend.text = element_text(size = 12, colour = "gray40"),
        strip.background =element_rect(colour = "gray40", fill = "honeydew2"),
        strip.text = element_text(size = 12, colour = "gray40"),
        legend.position = "right",
        legend.key = element_blank()) +
  guides(colour = guide_legend(title="Species", nrow = 2), shape = guide_legend(title="Dataset\nType", nrow = 2))
```
